# Matrix Vesicles from Osteoblasts Promote Atherosclerotic Calcification

**DOI:** 10.1101/2024.04.18.590180

**Authors:** Xiaoli Wang, Jie Ren, Zhen Zhang, Fei Fang, Erxiang Wang, Jianwei Li, Weihong He, Yang Shen, Xiaoheng Liu

## Abstract

**Backgrounds:** Vascular calcification often occurs with osteoporosis, a contradictory association known as “the calcification paradox”. Osteoblast-derived matrix vesicles (Ost-MVs) have been implicated in bone mineralization, and also have a potential role in ectopic vascular calcification. Herein, we aim to investigate the contributions that Ost-MVs make to the bone–vascular calcification paradox and the underlying mechanisms.

**Methods and Results:** Hyperlipidemia-induced atherosclerotic calcification in mice was accompanied with bone mineral loss, as evidenced by reduced deposition of Ost-MVs in the bone matrix and increased release of Ost-MVs into the circulation. Intravenous injection of fluorescent DiІ-labeled Ost-MVs revealed a marked fluorescence accumulation in the aorta of atherogenic mice, whereas no fluorescence signals were observed in normal controls. Using proteomics to analyze proteins in non-matrix bound Ost-MVs and mineralized SMC-derived MVs (SMC-MVs), we found Lamp1 was specifically expressed in SMC-MVs, and Nid2 was exclusively expressed in Ost-MVs. We further demonstrated that both Lamp1 and Nid2 were co-localized with Collagen І within calcific plaques, indicating the involvement of both Ost-MVs and SMC-MVs in atherosclerotic calcification. Mechanistically, LPS-induced vascular injury facilitated the transendothelial transport of Ost-MVs. The recruitment of circulating Ost-MVs was regulated by remodeled Collagen І during calcification progression. Furthermore, the phenotypic transition of SMCs determined the endocytosis of Ost-MVs. Finally, we demonstrated that either recruited Ost-MVs or resident SMC-MVs accelerated atherosclerotic calcification, depending on the Ras-Raf-ERK signaling.

**Conclusion:** Atherosclerotic calcification-induced Ost-MVs are released into circulation, facilitating the transport from bone to plaque lesions and exacerbating artery calcification progression. The mechanisms of Ost-MVs recruitment include vascular injury allowing transendothelial transport of Ost-MVs, collagen І remodeling promoting Ost-MVs aggregation, and SMC phenotypic switch to facilitate Ost-MVs uptake. Our results further revealed that both recruited Ost-MVs and calcifying SMC-MVs aggravate calcification through the Ras-Raf-ERK pathway.

## 1 Introduction

Cardiovascular disease (CVD) is a serious health challenge, causing more deaths world-wide than cancer.^1^ Atherosclerosis represents the culprit of cardiovascular morbidity and mortality. A typical pathologic manifestation of advanced atherosclerosis is intimal calcification, which is the ectopic deposition of apatite mineral in the intimal layers of the vessel wall. A growing number of studies have demonstrated that microcalcification in vulnerable plaques contributes to plaque destabilization and fatal plaque rupture.^2, 3^

Resembling the mineralization of bone, intimal calcification is an active process in which vascular smooth muscle cells (SMCs) undergo osteogenic transformation. This process is initiated by the deposition and aggregation of mineralization-competent matrix vesicles (MVs). ^4–6^ MVs, also called calcifying extracellular vesicles (EVs), are small membranous structures (30–300 nm in diameter), and are released by mineral-forming cells, including osteoblasts, chondrocytes, odontoblasts and calcifying SMCs.^7–9^ Functionally, MVs are thought to serve as nucleation sites for calcium crystal formation through mineralization-related components, including tissue non-specific alkaline phosphatase (TNAP), annexins (ANX; principally ANX2, 5 and 6) and phosphatidyl serine (PS).^10^ MVs that sequester Ca^2+^ and PO_4_^3−^ ions aggregate and deposit in extracellular matrix (ECM) by interaction with type І collagen, and the aggregation of MVs initiate the formation of microcalcifications and ultimately lead to macrocalcification.^11, 12^ Furthermore, MVs possess a metabolically active outer membrane that protects the internal cargo, consisting of proteins, microRNA, and other components from the parental cell. Emerging studies have suggested that MVs play a critical role in intercellular communication through their capacity to transfer their internal cargoes.^11–14^ Currently, despite an increased understanding of the role of MVs in atherosclerotic calcification, the heterogeneity of MV populations renders largely unknown regarding the cellular origins of MVs responsible for atherosclerotic calcification. Further investigation is warranted to elucidate the specific mechanisms of MV actions in the progression of ectopic arterial calcification.

Existing studies have provided evidence that mineralized SMCs located in the calcification niche produce MVs (SMC-MVs) to promote mineral nucleation, leading to the development of microcalcifications and ultimately large calcification.^9, 11, 12, 15^ SMCs, as the predominant cell type in the arterial vessel wall, play a crucial role in vascular calcification. Under physiological condition, SMCs adopt a quiescent state, often referred to contractile phenotype, and express SMC markers [e.g., α-smooth muscle actin (α-SMA), smooth muscle 22α (SM22α) and smooth muscle myosin heavy chain (SMMHC)]. And healthy SMCs efficiently prevent vascular calcification by expression of calcification inhibitors, some of which are loaded into MVs. In the context of pathological calcification, the change from the contractile to an osteochondrogenic phenotype is characterized by the development of MVs, down-regulation of mineralization inhibitory molecules, and elaboration of a calcification prone matrix.^5, 16^ This phenotype is accompanied by loss of SMC markers, and gain of osteochondrogenic markers [e.g., Runx2, osteopontin, osteocalcin, alkaline phosphatase (ALP) and collagen І].^5, 16^

More importantly, numerous epidemiological studies have provided evidence of a correlation between osteoporosis and cardiovascular disease.^17,18^ The progression of ectopic calcification is often accompanied by either decreased bone mineral density (BMD) or disturbed bone turnover.^19–21^ This contradictory association is known as “the calcification paradox”. However, the mechanism underlying this calcification paradox is not fully understood. Studies have indicated that multiple growth factors deposited in the bone matrix can be released into the bone marrow during active bone remodeling to regulate of bone homeostasis.^22, 23^ Moreover, evidence from clinical observations, animal models, and molecular studies indicates that factors such as BMP-2, MGP, OPN, OPG and inorganic pyrophosphate not only regulate bone cell differentiation and mineralization, but also play a role in vascular calcification.^24^ These finding, together with the well-described role of MVs as nucleation sites and intracellular messengers, suggest that osteoblast-derived MVs (Ost-MVs) may be released into the circulation via bone marrow to induce calcification paradox.^14, 25, 26^ Therefore, we hypothesized that Ost-MVs act as key mediators linking bone remodeling with ectopic vascular calcification and contribute to the progression of atherosclerotic calcification.

In this study, we elucidate the role of Ost-MVs in the bone–vascular calcification paradox within atherosclerotic calcification. We have demonstrated that Ost-MVs released from atherosclerotic calcification-induced bone remodeling are transported from bone to calcification niche within plaque lesions. The underlying mechanisms of Ost-MVs recruitment involve endothelial dysfunction facilitating Ost-MVs transendothelial transport, Collagen І remodeling-mediated Ost-MVs deposition, and SMC phenotypic switch to facilitate Ost-MVs uptake. Our results further reveal that both recruited Ost-MVs and resident SMC-MVs promote osteogenic differentiation of recipient SMCs and aggravated calcification through the Ras-Raf-ERK signaling pathway. Our work contributes to the body of evidence linking bone remodeling and arterial calcification, leading to a more comprehensive understanding of the pathobiological mechanisms underlying the calcification paradox and informing the development of new therapeutic strategies.

## 2 Methods

### Ethics statement

Animal experiments were approved by the Animal Ethics Committee and followed the Guidelines for the Care and Use of Laboratory Animals at Sichuan University (Sichuan China). Animals’ procedures were carried out in accordance with guidelines from Directive 2010/63/EU. Mice were housed in the Laboratory Animal Unit during experimental period. All mice were kept in an environment with a temperature of 22±1 °C, relative humidity of 50±1%, and a 12h light/dark cycle. For animal experiments, mice were anesthetized with 1% sodium pentobarbital (i.p. injection, 50mg/kg) and were euthanized by exsanguination.

### Mice

Mice were purchased from GemPharmatech^TM^. Atherosclerotic calcification models were established by using apolipoprotein E-deficient (ApoE^−/−^) mice (6 to 8-week-old, male) with a 20-week high-fat diet (HFD) containing 15% fat and 0.25% cholesterol (Laboratory Animal Center of Sichuan University, China). Age-matched wild-type (WT) C57/BL6J mice consuming a normal diet (ND) for 20 weeks served as controls. Mice were sacrificed to harvest aortas, tibias, and femurs for follow-up assays.

### Micro-CT analysis

The dissected femurs were fixed with 4% paraformaldehyde for 24 h at room temperature and subsequently scanned ex vivo using a micro-CT scanner (mCT50, Scanco Medical, Bassersdorf, Switzerland). Three-dimensional reconstructions of femurs were performed with a micro-CT system. The bone morphometric parameters, including bone mineral density (BMD), ratio of bone volume to tissue volume (BV/TV), trabecular number (Tb.N), trabecular thickness (Tb.Th) and trabecular separation (Tb.Sp), were analyzed. The operator who performed the scan analysis was blinded to the treatment assignment of the specimens.

### Cell culture and treatment

The mouse aortic vascular smooth muscle cells (MOVAS) and MC3T3 cells were obtained from ATCC. Both cells were maintained in Dulbecco’s modified Eagle’s medium (DMEM, Gibco) supplemented with 10% fetal bovine serum, and 1% penicillin-streptomycin (control condition: normal medium (NM)). For osteogenic phenotype induction, SMCs and MC3T3 cells were cultured for up to 14d in the presence of either control medium or osteogenic medium (OM) consisting of NM with the following additions:10 nM dexamethasone, 10 mM β-glycerolphosphate, and 100 mM L-ascorbate phosphate. Medium were changed every 3 to 4 days. For SMC phenotypic switching assays, heparin (100 μg/mL, MCE) and platelet derived growth factor subunit B (PDGF-BB, 20 ng/mL, PeproTech) were used to induce the contractile or synthetic phenotype of SMCs, as previously described.^27^

### MVs tracing analysis

For MV tracing assay, MVs were labeled with red fluorescent dye DiІ (Beyotime, China) specific for the MV bilayer. Briefly, purified MVs were incubated with DiІ (5 μM) at 37°C for 10 min. Following the 10 min incubation, stained MVs were ultracentrifugated at 300g (20 min), 3000g (20 min),10000g (30 min), and 100 000g (2h) to remove unincorporated dye. The same volume of PBS without MVs treated with the same procedures simultaneously was used as non-vesicle controls. The stained MV pellets were then resuspended in PBS, and the protein concentration of MV suspension was determined using a BCA protein assay kit.

For analysis of MV biodistribution, DiІ-labeled MVs or non-vesicle controls were intravenously injected into male ApoE^−/−^ mice consumed HFD for 12 weeks or WT mice on a ND for 12 weeks at a concentration of 100 μg/100 μL per mouse. After 12h, the mice were sacrificed and their organs (aorta, heart, liver, spleen, lung, kidney, pancreas and bone) were harvested for fluorescence imaging using an *in vivo* imaging system (IVIS, PerkinElmer).

After IVIS imaging, the aortic arch was snap-frozen in OCT compound and sectioned, followed by analysis of immunofluorescence by confocal microscope (Zeiss, Germany). Meanwhile, organs (heart, liver, spleen, lung, kidney, and pancreas) were fixed in 4% paraformaldehyde, then embedded in paraffin and cut into sections (5 μm) for HE staining. To further evaluate the role of collagen І in vascular-targeting effects of Ost-MVs, Halofuginone (RU-19110, MCE), a collagen І synesis inhibitor, was used to treat ApoE^−/−^ mice (6 to 8-week-old, male). ApoE^−/−^ mice were fed a HFD for 12 weeks in total, and mice were simultaneously intraperitoneally injected with Halofuginone (0.25 mg/kg) once every week for 12 weeks. The control group of ApoE^−/−^ mice was administered an equivalent volume of PBS alone via intraperitoneal injection. At the end of the treatment period, the animals were sacrificed and their aorta was dissected for fluorescent imaging using IVIS as described above.

### Proteomic analysis of MVs

Proteins from collected SMC-MVs and Ost-MVs were separated in a 4–20% sodium dodecyl sulfate polyacrylamide gel electrophoresis (SDS-PAGE) gel (Bio-Rad, USA), and subjected to consequent analysis by liquid chromatography with tandem mass spectrometry (LC-MS/MS). Data were then quantified by a label-free quantification (LFQ) approach and analyzed using Perseus software.

### Immunofluorescence

For staining, an Alexa Fluor^TM^ 488 Tyramide SuperBoost^TM^ kit (Invitrogen, USA) was used according to the manufacturer’s instructions. In brief, sections were heated in citrate at 100°C for 15 min for antigen retrieval, followed by incubation with Blocking Buffer for 60 min at RT to block peroxidase activity. Aortic sections (5 μm) were then blocked with 10% goat serum, and incubated with primary antibody Lamp1 (ab62562, Abcam) for 60 min at RT. After washes, sections were incubated with the poly-HRP-conjugated secondary antibody for 60 min, and then incubated with Alexa FluorTM 488 tyramide reagent solution. After 5 min, the reaction was stopped with Reaction Stop Reagent, and slides were washed. Nid2 (ab14513, Abcam) and Collagen І (14695-1-AP, Proteintech) protein were stained by repeating the above steps. Cell nuclei were marked with DAPI (Solarbio, China) for 15 min at RT prior to being cover slipped with antifade reagent. The entire process was conducted in the dark, and all washes were done with PBS. Fluorescent images were acquired using a confocal microscope (Zeiss, Germany).

### Immunohistochemistry

Immunohistochemistry staining was used to determine the expression of Lamp1 (ab62562, Abcam) and Nid2 (ab14513, Abcam) in the paraffin-embedded aortic sections. In brief, aortic sections (5 μm) were heated at 65°C in the oven for 1h, followed by dewaxing in xylenes and then rehydrating through an alcohol series (100%, 95%, 80%, 75%). Antigen retrieval of sections was induced by heat in citrate at 100°C for 15 min, and endogenous peroxidases activity was diminished by hydrogen peroxide incubation for 10 min. Sections were then blocked by 5% goat serum for 1h, followed by incubating with primary antibody overnight at 4°C. The next day, sections were rewarmed at RT for 30 min and then incubated with a secondary antibody (ZSGB-BIO, China) for 30 min at RT. Thereafter, a DAB kit (Solarbio, China) was used to detect positive staining. Before optical imaging, sections were counterstained with haematoxylin, dehydrated, and mounted.

### Transwell assays

Cells were co-cultured indirectly in a 24 transwell plates (Corning, USA). Human umbilical vein endothelial cells (HUVECs, 5 × 10^5^ cells/mL) were cultured in the upper chamber of the Transwell insert, and human aortic smooth muscle cells (HASMCs, 1 × 10^5^ cells/well) were cultured in the lower chamber (in the 24-well plate). Before co-culture with SMCs, HUVECs were pretreated with 1 μg/mL LPS (L2880, Sigma) for 48h. The control group was treated with an equal volume of DMEM. After LPS treatment, the Transwell inserts were transferred to the wells seeded with the HASMCs, and the culture medium was changed into conditioned NM (0.1% FBS). Dil-labeled Ost-MVs (15 μg) were then added to the upper chamber, and continued to be incubated for 12h. Thereafter, HASMCs were washed with PBS and fixed in 4% paraformaldehyde for 30 min. After washes, the cells were incubated with FITC-labeled phalloidin(Solarbio, China) for 30 min, followed by DAPI staining for 15 min. Before imaging, cells were washed, air-dried, and cover-slipped with antifade reagent.

### Collagen hydrogel experiments

Collagen hydrogels with different concentrations (0.1 mg/mL, 0.5 mg/mL, 1 mg/mL and 2 mg/mL) were made by slowly adjusting the pH to 7 of high concentration rat tail collagen I (Solarbio, China) stored in a solution of acetic acid. The neutralized collagen solution was added to a sterile round glass slide (1 cm in diameter) within a 24-well plate overnight at 37 °C. The undried collagen solution was discarded, and slides were gently rinsed with PBS. MVs (15 μg) or DiІ-labeled MVs (15 μg) in conditioned OM (0.1% FBS) were then added to the collagen hydrogels and incubated for 3 days at 37°C. The accumulation and aggregation of MVs on collagen hydrogels were visualized by confocal microscope (Zeiss, Germany) and SEM (ThermoFisher, USA), respectively. For fluorescent imaging, hydrogels were fixed in 4% paraformaldehyde, and blocked with BSA (1%) for 30 min. Samples were then incubated with primary antibody collagen І (14695-1-AP, Proteintech) at 4 °C overnight, followed by incubation with a secondary antibody (ZSGB-BIO, China) for 60 min at RT. After washes, slides were cover-slipped with antifade reagent, and imaged with a confocal microscope. For SEM observation, after fixation in 4% glutaraldehyde and washes with PBS, slides were dehydrated in an ascending alcohol series (30%, 50%, 70%, 90%, and 100%). The samples were then mounted on stubs, vacuum-dried, and coated with a gold film before observation by SEM.

### MVs Uptake

MOVAS (1×10^5^ cells/mL) were seeded onto a sterile round glass slide (1 cm in diameter) within a 24-well plate and cultured with NM overnight at 37°C. Meanwhile, purified SMC-MVs or Ost-MVs were incubated with 5 μM red fluorescent dye DiІ (Beyotime, China) or green fluorescent dye DiO (Beyotime, China) for 10 min at 37°C, respectively. The unbound dye was removed by another round of MV isolation procedures as described above. Stained MV pellets were then resuspended in PBS, and the protein concentration of MV suspension was determined using a BCA protein assay kit. Before co-culture, change NM into conditioned OM (0.1% FBS). Then, Dil-labeled SMC-MVs (15 μg) and DiO-labeled Ost-MVs (15 μg) were added to SMCs and incubated for 12h, 24h and 48h at 37°C, 5%CO_2_. For confocal imaging, cells were washed and fixed in 4% paraformaldehyde. The nuclei were stained with DAPI at RT. After the final washes, slides were cover-slipped with antifade reagent, and observed with a confocal microscope (Zeiss, Germany).

### Flow cytometry analysis

MOVAS were grown in a T25 culture flask to induce contractile or synthetic phenotype as previously described. Dil-labeled SMC-MVs (50 μg) and DiO-labeled Ost-MVs (50 μg) were then added to contractile/synthetic SMCs, and incubated for 12h at 37°C, 5%CO_2_. After incubation, SMCs were rinsed with PBS and underwent trypsin digestion for preparation of a single cell suspension. The resulting cell suspension was centrifuged at 1000 rpm for 3 min, and resuspended in PBS at a concentration of 1×10^6^ cells/mL. The uptake of SMC-MVs and Ost-MVs by SMCs was analyzed with flow cytometry (BD, FACS Celesta).

### In vitro mineralization assays and ALP activity

MOVAS were incubated with/without SMC-MVs/Ost-MVs (50 μg) in the presence of procalcifying OM (to provide phosphorus, which is needed for calcification) for up to 7 days. For alizarin red staining, cells were fixed in 95% ethanol and stained with 1%, PH4.2 alizarin red s (Solarbio, China) for 30 min at RT. Excess dye was removed by washing with distilled water, and images were acquired with an optical microscope. To quantify alizarin red staining, we added a 100 mmol/L hexadecyl-pyridinium chloride (C9002, Sigma) solution to SMCs for 30 min at 37°C. The optical density (OD) was measured with a spectrophotometer at 560 nm using hexadecyl-pyridinium chloride solution as a blank. For ALP activity analysis, MVs-treated SMCs was measured using the Alkaline Phosphatase Activity Colorimetric Assay Kit (Beyotime, China) according to the manufacturer’s instructions. The absorbance was measured using a microplate reader at a wavelength of 405 nm.

### Transcriptome analysis

After incubation with/without SMC-MVs/Ost-MVs (50 μg) in the presence of pro-calcifying OM for up to 7 days, MOVAS were washed and collected for subsequent transcriptome sequencing (RNA-seq).

### Western blotting

MV or cell samples were lysed with RIPA buffer containing protease and phosphatase inhibitors. Protein concentration was measured using a BCA protein assay kit (Beyotime, China). Western blotting was performed with the use of anti-ANX2 antibody (60051-1-lg, Proteintech), anti-ANX5 antibody (ab14196, Abcam), anti-ANX6 antibody (12542-1-AP, Proteintech), anti-Lamp1 antibody (ab62562, Abcam), anti-Nid2 antibody (ab14513, Abcam), anti-Collagen І antibody (66761-1-Ig, Proteintech), anti-Ras antibody (ET1601-16,HuaBio), anti-Raf antibody (ET1701-21, HuaBio), anti-ERK antibody (4695, CST), anti-p-ERK (4370, CST), anti-α-SMA antibody (ab7817, Abcam), anti-SM22α antibody (ab14106, Abcam), anti-Collagen І antibody (14695-1-AP, Proteintech), anti-ALP (ET1601-21, HuaBio), and anti-GAPDH antibody (21612, SAB) overnight at 4°C, followed by the HRP-labeled anti-rabbit or anti-mouse IgG secondary antibodies (ZSGB-BIO, China.). Protein expression was detected and analyzed using an ECL substrate reagent kit (4A Biotech, China) and Molecular Image®ChemiDocTM XRS+ system with Image Lab^TM^ Software.

### In vivo mineralization assays

To assay the development of atherosclerotic calcification in SMC-MVs and Ost-MVs-treated ApoE^−/−^ mice and its regulation by MAPK signaling, six-week-old male ApoE^−/−^ mice were randomly assigned into five groups: the control group (HFD), the SMC-MVs group (HFD+SMC-MVs), the Ost-MVs group (HFD+Ost-MVs), the SMC-MVs+PD98059 group (HFD+SMC-MVs+PD98059), and the Ost-MVs+PD98059 (HFD+Ost-MVs+PD98059), with 5 mice in each group. SMC-MVs or Ost-MVs were injected intravenously for 12 weeks (100 μg/100 μL per mouse). Simultaneously, mice were treated with 0.3 mg/kg PD98059 (A1663, APExBIO). The control group was administered an equivalent volume of PBS. Both MVs and PD98059 administrations were performed every week. After 12 weeks, the aorta was excised and fixed in 4% paraformaldehyde, then embedded in paraffin and cut into sections (5 μm) for calcific deposits determination. Calcific deposits in aortas were visualized by Alizarin Red S staining. Briefly, sections were dewaxed in xylene and rehydrated in alcohol series. Then, sections were washed with PBS and incubated with 1%, PH4.2 Alizarin Red S (Solarbio, China) for 30 min at 37°C. Histological staining images were obtained using an optical microscope.

### Statistical analysis

Data are shown as mean ± standard deviation (SD), and n indicates the number of biological samples followed by the number of independent experiments. For comparison between two groups, an unpaired two-tailed Student’s t test was performed. For comparison among three or more treatment groups, one-way ANOVA with Tukey’s post hoc, and two-way ANOVA with Sidak correction were done. Significance was considered when P<0.05.

## 3 Results

### 3.1 Atherosclerotic calcification was accompanied by Ost-MVs-mediated bone loss

To explore the relationship between artery calcification and bone metabolism, we developed a hyperlipidemia-induced atherosclerotic calcification model using ApoE^−/−^mice fed a high-fat diet. Then, the formation of calcification and changes in bone mineral density (BMD) were determined. In the arch of the aorta, fibro-fatty nodules in the advanced atherosclerotic lesions were a nidus for calcification and became highly calcified in ApoE^−/−^ mice consumed a HFD for 20 weeks (Figure 1A). And the osteogenic transformation of SMCs was characterized by an elevated expression of osteogenic makers (ALP and RUNX2) in HFD-treated ApoE^−/−^ mice (Supplementary material online, Figure S1A-B). As viewed under TEM, MVs bound to collagen in acellular ECM regions within calcified areas of atherosclerotic plaques (Figure 1B). NTA measurements further revealed that the average diameter of isolated MVs from the vascular matrix was in anticipated size range (30-300 nm) (Figure 1C). More importantly, there was an increase in particle concentration of MVs in calcified mice compared with that in the control mice (normal diet–fed C57/BL6J mice) (Figure 1D). Extensive researches have been conducted on the significant accumulation of MVs during atherosclerotic calcification, particularly in relation to increased secretion of resident mineralized SMCs,^11, 15, 28, 29^ and the release of circulating MVs from aged bone.^30^

**Figure 1.**
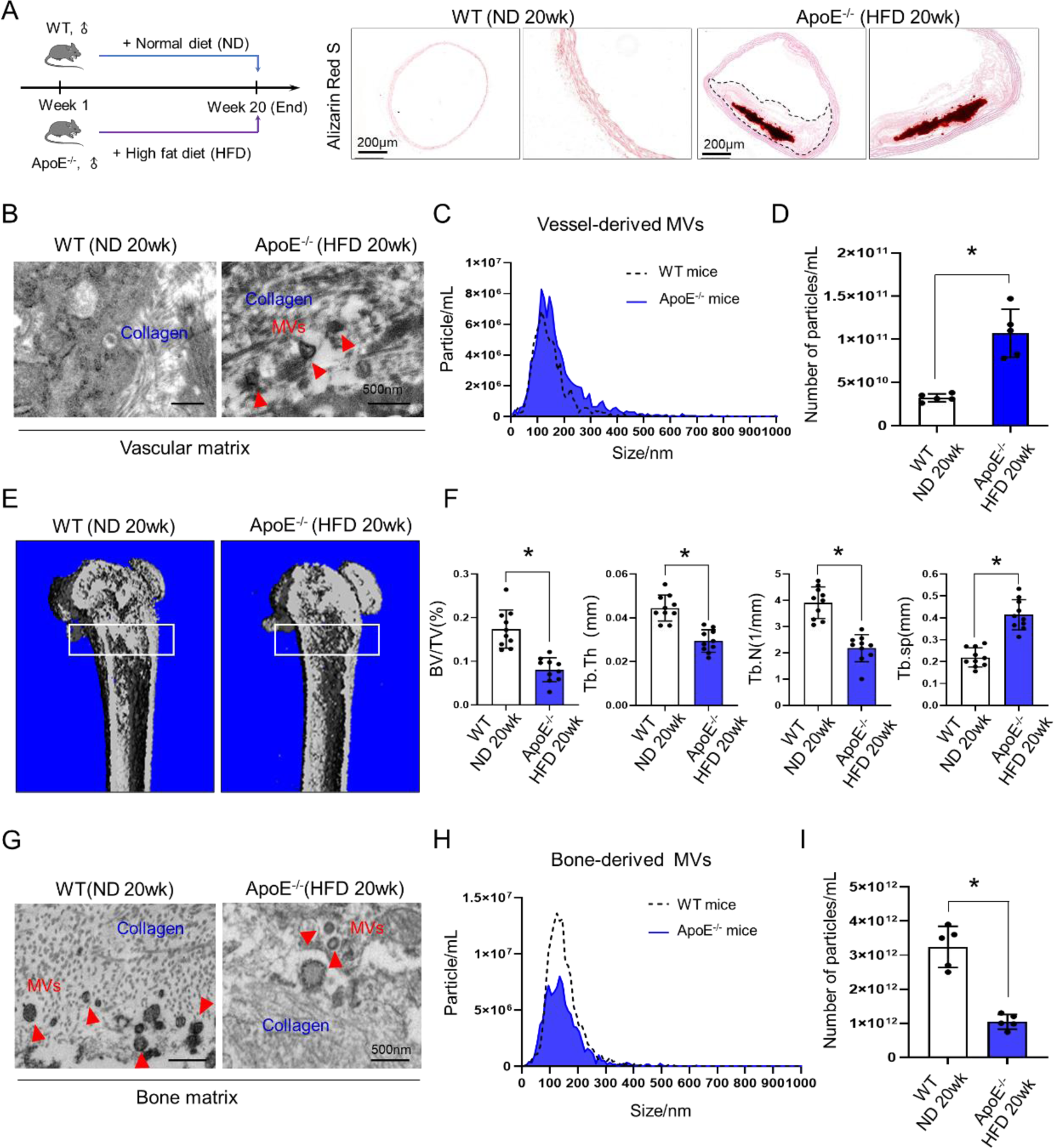
Atherosclerotic calcification was accompanied with Ost-MVs disorder-induced bone loss. (A) Atherosclerotic calcification in HFD-induced ApoE^−/−^ mice. Representative images of aortic sections stained by Alizarin Red S (red) showed significantly increased calcium deposition in plaque lesions of ApoE^−/−^ mice consumed HFD for 20 weeks compared to that of ND-treated WT mice (n=10), scale bar=200 μm. (B) TEM images of deposited MVs (red arrows indicate MVs with membrane hydroxyapatite formation) in vascular matrix of normal and calcified vessels (n=5), scale bars= 500 nm. (C) The size distribution of vascular matrix-derived MVs measured by NTA. (D) Statistical analysis of the number of vascular matrix-derived MVs in normal mice and calcified mice (n=5). (E) Trabecular bone was assessed by microCT. Representative bone cross sections from WT mice on a chow diet for 20 weeks and ApoE^−/−^ mice consumed HFD for 20 weeks (n=10). (F) Bone volume/tissue volume (BV/TV), trabecular thickness (Tb.Th), trabecular number (Tb.N), and trabecular separation (Tb.Sp). (G) TEM images showed accumulation of MVs (red arrows indicate MVs with membrane hydroxyapatite formation) in the bone matrix of normal and calcified mice (n=10). Scale bars= 500 nm. (H) Size histograms of bone-derived MV particles obtained by NTA measurement. (I) Statistical analysis of the number of bone-derived MVs in normal control mice and calcified mice (n=5). *, P<0.05, the difference between groups was statistically significant.

We next assessed the bone morphology of calcified ApoE^−/−^ mice and normal controls. In contrast to controls, microCT analysis showed a decrease in bone mass at the cancellous region of femur of calcified mice. This was characterized by a significant reductions in the ratio of bone volume to tissue volume (BV/TV), trabecular thickness (Tb.Th), and trabecular number (Tb.N), and an increase in the trabecular separation (Tb.Sp) (Figure 1E-F). Moreover, the deposited MVs in bone matrix of normal and calcified mice were visualized by TEM. In contrast to abundantly deposited MVs in ECM of normal controls, a significant reduction in the matrix deposition of MVs was observed in calcified mice (Figure 1G). Further quantitative analysis was performed by isolation of MVs from the tibias and femurs of normal and calcified mice. As shown in Figure 1H, the average diameter of isolated MVs was 128.7±3.6 nm in control group, and 144.3±1.3 nm in calcific group, consistent with the expectation of a 30-300 nm size range. And quantitative analysis suggested a decrease in particle concentration of MVs in mice with calcification compared with normal controls (Figure 1I).

Meanwhile, we analyzed the changes in the production of Ost-MVs in vitro (Supplementary material online, Figure S2A). Osteoblasts were isolated from control and calcified mice, and then matrix-bound MVs and free medium MVs were collected with a serial centrifugation method. TEM imaging and NTA analysis indicated that Ost-MVs presented a typical bi-layered, cup-like morphology in the size range of 30-300 nm (Supplementary material online, Figure S2B, D and F). Results of western blots showed that ANX2, ANX5 and ANX6 were highly enriched in Ost-MVs, further confirming the MVs identity of these vesicles (Supplementary material online, Figure S2C). Likewise, we found the concentration of Ost-MVs in the medium was significantly increased, while the level of matrix-bound Ost-MVs was reduced in the calcific group (Supplementary material online, Figure S2E and J).

The combined evidence from in vitro and in vivo studies suggests a close relationship between atherosclerotic calcification and bone loss, with Ost-MVs released into the circulation appearing to be responsible for the bone–vascular calcification association.

### 3.2 Ost-MVs releasing into circulation targeted atherosclerotic lesions

To track in vivo biodistribution of releasing Ost-MVs in the setting of atherosclerosis, we first prepared Ost-MVs by culturing preosteoblast-like cells, MC3T3 cells (osteoblast precursor cell line), with calcifying osteogenic medium (OM); by contrast, cells in control group were under the condition of normal medium (NM). NTA and TEM results showed that OM-induced Ost-MVs were small, 30-300nm diameter, membrane-bound vesicles (Figure 2A-B), similar in size and morphology to previously described MVs.^10, 31^ The vesicle identity of isolated Ost-MVs was further evidenced by the expression of surface markers ANX2, ANX5 and ANX6, which were highly enriched in OM-induced Ost-MVs compared with that of NM-treated group (Figure 2C).

**Figure 2.**
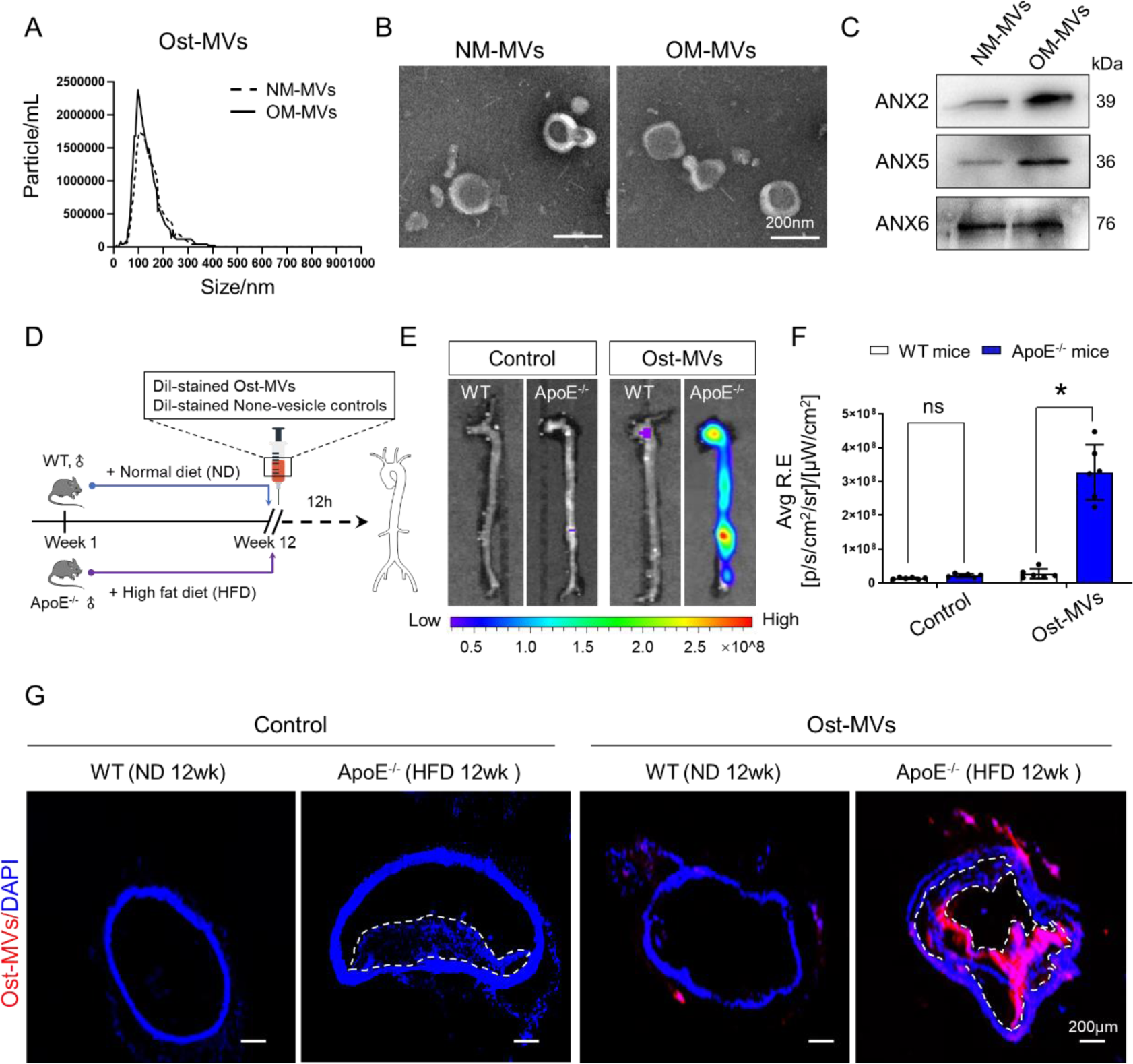
Circulating Ost-MVs targeted plaque lesions in the context of atherosclerosis. (A-C) Identification of Ost-MVs released by MC3T3 cells cultured in NM or OM. NTA (A) and TEM (B) were used to measure, quantify and visualize Ost-MVs. Scale bars= 200 nm. (C) Western blot revealed the enrichment of ANX2, ANX5, and ANX6 in mineralization-competent MVs. (D) Experimental design of in vivo assay to track the distribution of the fluorescence signal in mice after injection of after injection of non-vesicular controls or Dil-labeled Ost-MVs. (E) Biophotonic images of the arterial distribution of the fluorescence signal in aortas (n=6). (F) Statistical analysis of the average fluorescence intensities in arteries (n=6). *, P<0.05, the difference between groups was statistically significant. (G) Confocal microscopy images of the fluorescent signal distribution in aortic sections of mice after injection of non-vesicular controls or Dil-labeled Ost-MVs (n=6). Scale bars= 200 μm.

Then, Ost-MVs were labeled with a red fluorescent dye DiІ and subsequently injected into ND-treated controls and HFD-treatment ApoE^−/−^ mice via the tail vein for 12h, respectively (Figure 2D). Following Ost-MV administration, the aortas were harvested and ex vivo analysis of the distribution of stained Ost-MVs was assessed using IVIS according to previous reported studies.^32–35^ And the amount of Ost-MVs accumulated to the aorta was measured through fluorescent intensity. As shown in Figure 2E-F, we observed a strong fluorescence signal in the aortas of ApoE^−/−^ mice, in particular the athero-susceptible regions such as the aortic arch and abdominal aorta, while there was almost no fluorescence signal in aortas of control mice. Similar results were obtained by immunofluorescence, which showed that Ost-MVs were localized and accumulated within the atherosclerotic plaque (Figure 2G). These findings provide further evidence that Ost-MVs entering circulation during atherogenesis-related bone remodeling could be transported to blood vessels.

In addition, we measured the accumulation of stained Ost-MVs in other organs after intravenous administration, including heart, liver, spleen, lung, etc. As shown in Supplementary material online, Figure S3A-B, Ost-MVs accumulated mainly in the liver and lungs. We also evaluated the biocompatibility of Ost-MVs in vivo after intravenous injection of Ost-MVs or non-vesicle controls. There were no apparent histomorphometric changes in the heart, liver, spleen, lung, kidney, and pancreas with Ost-MVs injection, suggesting that Ost-MVs were well-tolerated, with little to no acute, systemic toxicity (Supplementary material online, Figure S3C).

### 3.3 Both circulating Ost-MVs and resident SMC-MVs contributed to the formation of atherosclerotic calcification

Extensive studies have evidenced that calcific mineral formation and maturation in atherosclerotic plaques results from a series of events involving the aggregation of calcifying SMC-MVs, the formation of microcalcifications and ultimately large calcification zones.^11, 15, 28, 29^ As shown in Figure 2, we tracked that circulating Ost-MVs could naturally target atherosclerotic lesions. To further elucidate the cellular origins of MVs that are mainly responsible for atherosclerotic calcification, we first performed label-free quantitative proteomics to assess the protein profiles of Ost-MVs and SMC-MVs isolated from calcification-promoting OM. To do this, OM-induced medium SMC-MVs were collected and characterized by three distinct methods (NTA, TEM and western blot). NTA results showed that both Ost-MVs and SMC-MVs were 30 to 300nm in diameter (Figure 2A, and see Supplementary material online, Figure S4A). TEM imaging and western blotting revealed Ost-MVs and SMC-MVs distinguishable with lipid bilayers, and showed abundances in MV marker proteins (ANX2/5/6) (Figure 2B-C, and see Supplementary material online, Figure S4A-B). Moreover, SEM with energy dispersive X-ray microanalysis demonstrated significantly elevated calcium:phosphorus ratio (Ca/P) in OM-induced Ost-MVs (1.39 vs. 2.26) and SMC-MVs (1.02 vs. 2.13) compared with corresponding control NM groups (Supplementary material online, Figure S4C). These results together validated that purified Ost-MVs and SMC-MVs were used for proteomics.

Next, using mass spectrometry, we identified 619 proteins in OM-induced Ost-MVs and SMC-MVs. As shown in Figure 3A-B, of 619 detected proteins, 184 were increased and 9 were decreased in SMC-MVs compared with Ost-MVs. The biological processes analysis showing enrichment of different proteins in two types of MVs found that the most abundant proteins were involved in cell-substrate adhesion, while the cellular component analysis identified that enriched proteins mainly originated from membrane region. Protein enrichment based on molecular functions indicated that most proteins of SMC-MVs showing significant differences were closely associated with cell adhesion molecule binding (Figure 3C). More importantly, through proteomics methods, we identified specific proteins such as Itga5 (Integrin alpha-5), Lamp1 (Lysosome associated membrane protein 1), Rab10 (Ras-related protein Rab-10), Clta (Clathrin light chain A), M6pr (Cation-dependent mannose-6-phosphate receptor), Cltb (Clathrin light chain B), Lama5 (Laminin subunit alpha-5), Itga7 (Integrin alpha-7), Scarb2 (Lysosome membrane protein 2), Itga3 (Integrin alpha-3) and Col4a2 (Collagen alpha-2(IV) chain) exclusively present in SMC-MVs, while Clu (Clusterin) and Nid2 (Nidogen 2) proteins were solely detected in Ost-MVs (Figure 3D). Results of western blotting confirmed that Lamp1 was specifically present in SMC-MVs, and Nid2 was expressed in Ost-MVs only (Figure 3E). Accordingly, Lamp1 and Nid2 were suggested as specific biomarkers of SMC-MVs and Ost-MVs in the present study.

**Figure 3.**
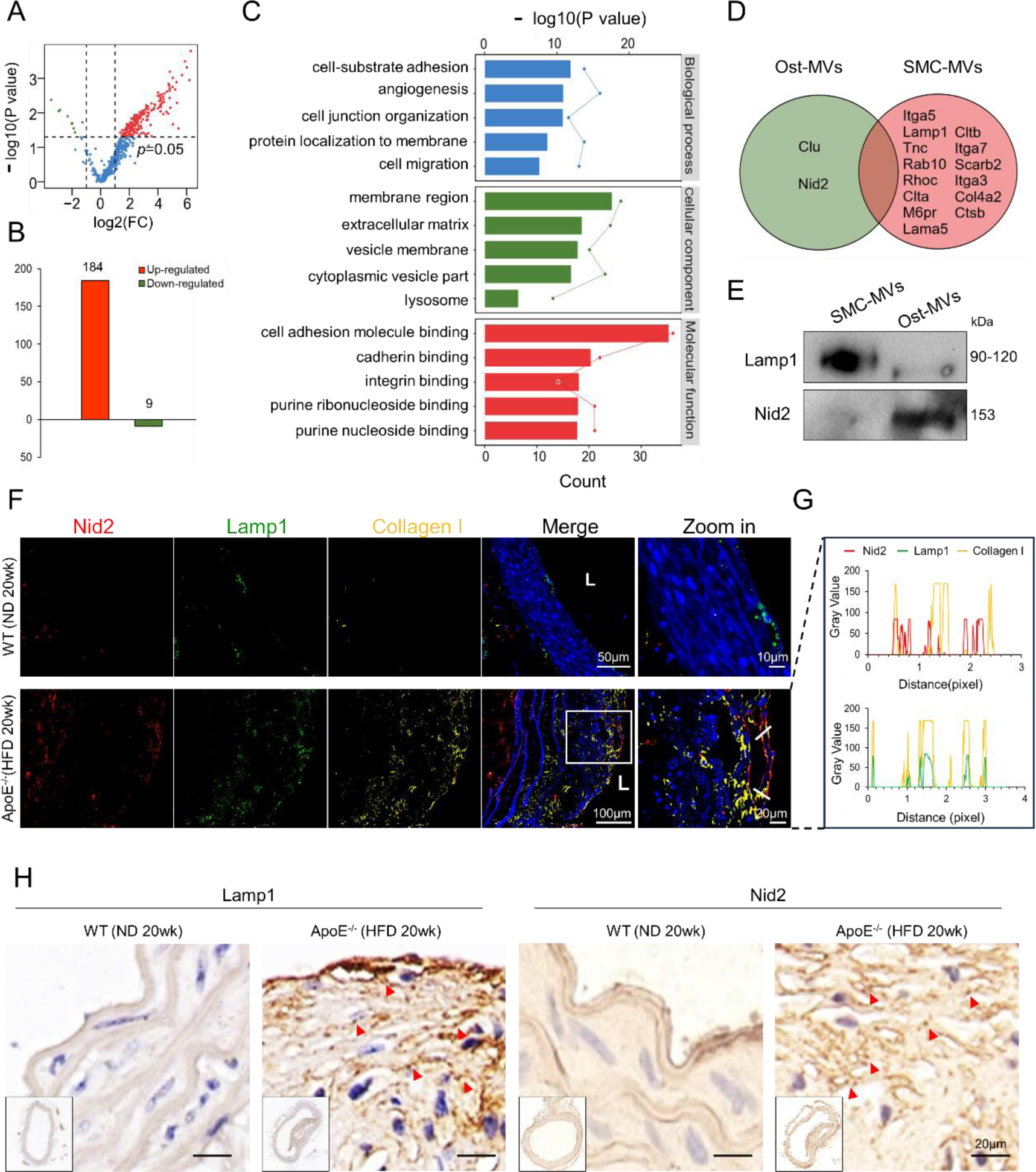
Both recruited Ost-MVs and resident SMC-MVs contributed to atherosclerotic calcification. (A) Proteomics protein volcano plot analysis for Ost-MVs and SMC-MVs isolated from pro-calcifying medium. (B) Histogram showed increased and decreased SMC-MVs protein abundances compared with Ost-MVs. (C) Functional enrichment of differentially expressed proteins in SMC-MVs compared with Ost-MVs. Bar plots represented the gene ontology enrichment analysis of differentially expressed proteins. (D) Venn diagram of detected proteins in Ost-MVs and SMC-MVs, with the green circle including proteins specifically detected in Ost-MVs and the red circle including proteins specifically detected in SMC-MVs. (E) Western blotting was used to determine the expression of Lamp1 and Nid2 in Ost-MVs and SMC-MVs. (F) Immunofluorescence staining of Nid2 (red), Lamp1 (green), and Collagen І (yellow) in aortic sections from control and calcified mice (n=5). Scale bars=50 and 100 µm. Designated regions indicated by white square frames are enlarged to show the details. Scale bars=10 and 20 µm. (G) Immunofluorescence co-localization analysis of Nid2/Lamp1 with Collagen І in calcific arteries. (H) Immunohistochemical analysis of Lamp1 and Nid2 expression in artery samples from control and calcifying ApoE^−/−^ mice donors (n=5). Arrows indicate extracellular, punctate Lamp1 or Nid2 staining. Scale bars=20 µm.

Finally, based on screened biomarkers of SMC-MVs and Ost-MVs, immunofluorescence and immunohistochemistry were used to determine the cellular origins of deposited MVs in calcified plaques. As shown in Figure 3F-G, both Lamp1 and Nid2 staining were detectable and co-localized with ECM protein collagen І in calcific plaques. Results of immunohistochemistry also indicated extracellular, punctate deposits of Lamp1 and Nid2 staining within plaques (Figure 3H), similar to previous observations of MVs in the ECM.^36^

Depending on specific markers in different origins of MVs, our results showed that circulating Ost-MVs, together with resident calcifying SMC-MVs, mediate the formation and development of atherosclerotic calcification.

### 3.4 Endothelial dysfunction and remodeled Collagen І during atherosclerosis directed the recruitment of Ost-MVs to vascular matrix

To explore the underlying mechanism by which circulating Ost-MV were recruited into the vascular matrix, we first examined whether Ost-MVs could move across endothelium under physiological and atherosclerotic conditions. Transwell models enabling the co-culture of LPS-pretreated endothelial cells (ECs) and SMCs were used to simulate healthy and impaired endothelium (Figure 4A). DiІ-labeled Ost-MVs were added to the upper chamber of transwell plates, and continued to culture for 12h. LPS-induced endothelial dysfunction markedly promoted the transendothelial transport of Ost-MVs and their uptake by SMCs grown on the bottom of the well, compared to the control group without LPS stimulation (Figure 4B-C).

**Figure 4.**
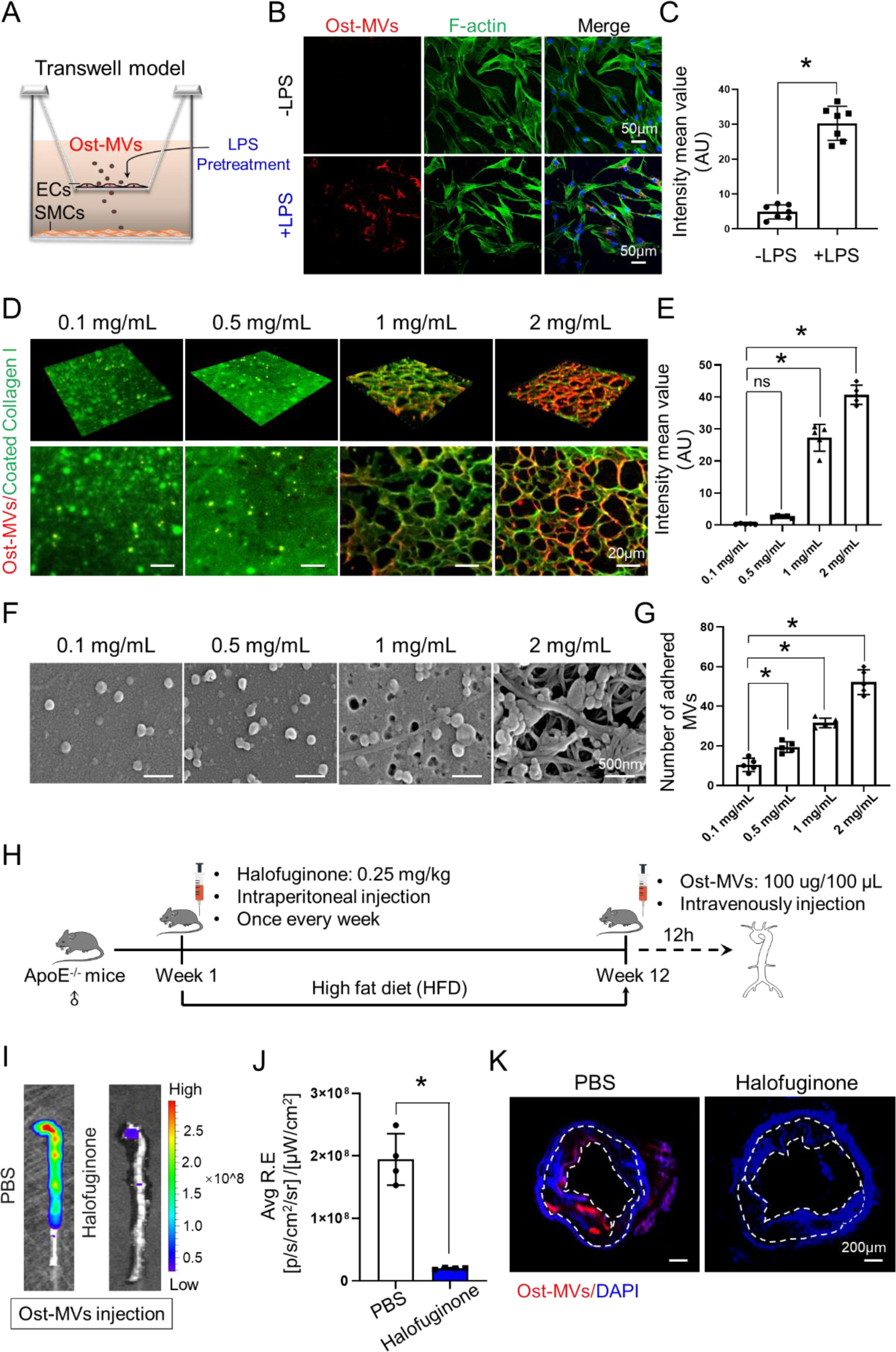
Endothelial injury and Collagen І remodeling during atherosclerotic calcification enabled the recruitment of circulating Ost-MVs. (A) Schematic diagram of the Ost-MVs transwell assay. ECs seeded on the top of the transwell chamber were pretreated with LPS for 48h, followed by co-culture with SMCs seeded on the lower chamber. Dil-stained Ost-MVs were then added to the upper chamber for 12h to explore whether Ost-MVs could transport across endothelium under physiological and pathological conditions. (B) Confocal images showed the transendothelial transport of Ost-MVs and the uptake of Ost-MVs by SMCs in the lower chambers. Red: Dil-stained Ost-MVs, Green: F-actin, Blue: DAPI. Scale bars=50 µm. (C) Statistical analysis of the average fluorescence intensities of DiI-labeled Ost-MVs in the lower chambers. (D) Confocal fluorescence images of MVs binding to collagen hydrogels following the introduction of DiІ-labeled Ost-MVs in calcifying medium for 3 days. The upper panel of fluorescence images showed 3D reconstruction from confocal z-stack images of collagen І (green) and Ost-MVs(red). Scale bars=20 µm. (E) Statistical analysis of the average fluorescence intensities of DiI-labeled Ost-MVs that were adhered and aggregated to the collagen hydrogels. (F) SEM images of MVs binding to collagen hydrogels following the introduction of DiІ-labeled Ost-MVs in calcifying medium for 3 days. The upper panel of fluorescence images showed 3D reconstruction from confocal z-stack images of collagen І (green) and Ost-MVs(red). Scale bars= 500 nm. (G) Statistical analysis of the number of adherent Ost-MVs binding to collagen hydrogels. (H) Experimental design of in vivo assay to test the role of collagen І in the recruitment of circulating Ost-MVs to the vascular matrix. ApoE^−/−^ (6 to 8-week-old, male) mice were fed a HFD for 12 weeks, and the mice underwent intraperitoneal injection of Halofuginone once every week simultaneously. Mice were then injected with Dil-labeled Ost-MVs at week 12 and harvested for sequent experiments. (I) Biophotonic images of the arterial distribution of the fluorescence signals in mice after injection of Dil-labeled Ost-MVs (n=4). (J) Statistical analysis of the average fluorescence intensities in arteries (n=4). (K) Representative images of the fluorescent signal distribution in aortic sections of mice after injection of Dil-labeled Ost-MVs. Scale bars= 200 μm. *, P<0.05, the difference between groups was statistically significant.

Besides, pathologic calcification and bone mineralization share several histopathological features, especially the secretion of large amounts of matrix components consisting largely of type І collagen, which acts as a scaffold to guide the aggregation of MVs, thereby promoting calcific mineral formation and maturation.^11, 37^ Hence, we speculated that the recruitment of circulating Ost-MVs to the vascular matrix may be ascribed to the shared feature of collagen І remodeling in both bone formation and vascular calcification. To test this hypothesis, we initially investigated the changes in collagen І expression during the progression of atherosclerotic calcification in vitro and in vivo. Consistent with previous studies, there was a significant increase in collagen І expression during SMC mineralization (Supplementary material online, Figure S5A-C). Then, we developed collagen hydrogels with different concentrations to mimic the alternation of collagen І at different stages of atherosclerosis. The addition of red fluorescently labeled Ost-MVs to the collagen hydrogels under pro-calcifying conditions allowed monitoring the binding of MVs to collagen І over time. At day 3, confocal fluorescence images revealed that DiІ-labeled Ost-MVs bound to collagen fibrils in a concentration-dependent manner (Figure 4D-E). Consistently, an increased number of Ost-MVs binding to collagen scaffold was observed by SEM with the increase of collagen І concentrations (Figure 4F-G).

Furthermore, we investigated the role of collagen І in the regulation of Ost-MVs recruitment to the vascular matrix in vivo. To achieve this, Halofuginone, a specific inhibitor of type I collagen synthesis, was selected.^38, 39^ The treatment with Halofuginone effectively inhibited the expression of collagen I in OM-treated SMCs (Supplementary material online, Figure S5D-E). In vivo, ApoE^−/−^ mice were fed an atherogenic diet for 12 weeks, and ApoE^−/−^ mice were injected with Halofuginone or PBS intraperitoneally once every week for 12 weeks. As previously described, DiІ-labeled Ost-MVs were then administrated via the tail vein of ApoE^−/−^ mice. After 12h from injection, the accumulation of stained Ost-MVs in aortas of ApoE^−/−^ mice was visualized using IVIS (Figure 4H). Our results found that Halofuginone treatment led to a significantly reduced fluorescent intensity in aortas, compared with PBS treatment (Figure 4I-J). Confocal microscopic images also showed that the aggregation of Ost-MVs within atherosclerotic plaque were remarkably inhibited by Halofuginone (Figure 4K). These findings suggest that vascular injury in atherosclerosis facilitates the transendothelial transport of circulating Ost-MVs, and that pathological collagen І alteration is essential for the recruitment of Ost-MVs to the vascular matrix.

### 3.5 Vascular smooth muscle cell phenotypic switching promoted recruited Ost-MVs uptake

Apart from the well-recognized role of nucleation sites, another emerging characteristic of MVs is their function as intracellular messengers by transferring their abundant functional cargoes.^30, 40, 41^ SMCs are the major cell type involved in the formation of calcification in atherosclerosis.^5^ To study the response of SMCs residing in plaque lesions to recruited Ost-MVs, we incubated SMC-/Ost-MVs with SMCs under pro-osteogenic conditions to simulate the developmental process of atherosclerotic calcification. SMC-MVs and Ost-MVs were labeled by fluorescent dyes DiІ (red) and DiO (green), respectively, and then the stained MVs were added into recipient SMCs. Interestingly, the result showed that SMCs were more prone to endocytose SMC-MVs at 12h and 24h, while the number of internalized Ost-MVs was significantly increased at 48h, suggesting a temporal difference in MVs uptake (Figure 5A-B).

**Figure 5.**
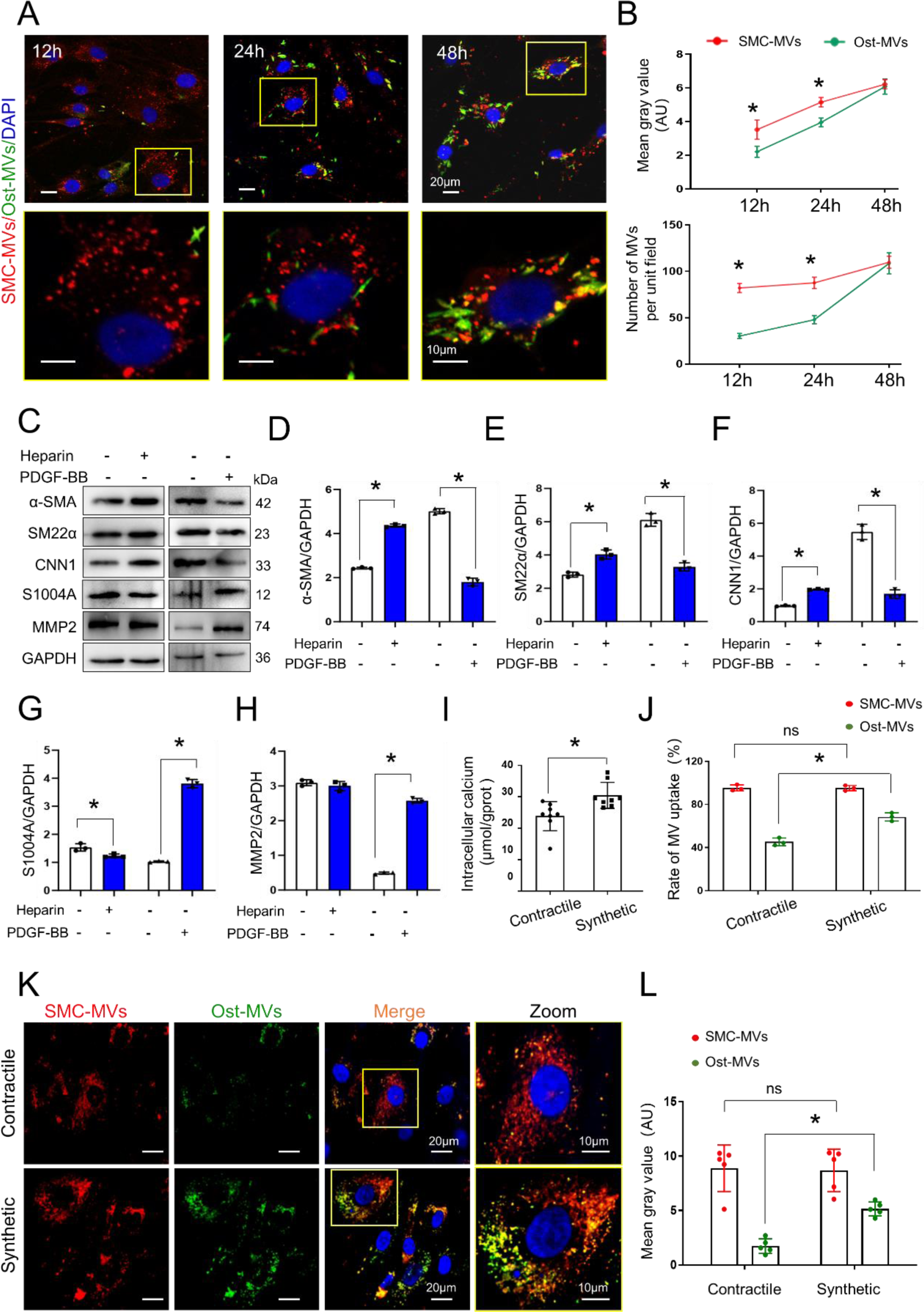
SMC phenotypic switching facilitated the endocytosis of Ost-MVs. (A) The uptake of SMC-MVs and Ost-MVs by recipient SMCs. DiІ-labeled SMC-MVs (red) and DiO-labeled Ost-MVs (green) were co-cultured with SMCs for 12h, 24h and 48h, and imaged by confocal microscopy. Scale bar=20 μm. (B) The fluorescence intensity of endocytosed SMC-MVs and Ost-MVs was statistically analyzed. (C-H) SMCs were treated with heparin (100 μg/mL) for 5d or PDGF-BB (20 ng/mL) for 2d to induce contractile and synthetic phenotype of SMCs, respectively. Western blotting and quantification of contractile makers (α-SMA, SM22α and CNN1) and synthetic markers (S1004A and MMP2) of SMCs. (I) Effects of SMC phenotypes on intracellular calcium content under OM condition. (J) Flow cytometric analysis of the uptake of SMC-MVs and Ost-MVs by contractile/synthetic SMCs. (K) Immunofluorescence assay was used to detect the internalization of DiІ-labeled SMC-MVs (red) and DiO-labeled Ost-MVs (green) by contractile/synthetic SMCs. Scale bar=20 μm. (L) The fluorescence intensity of endocytosed SMC-MVs and Ost-MVs by contractile and synthetic SMCs was statistically analyzed. Designated regions indicated by yellow square frames are enlarged to show the details. Scale bar=10 μm. *, P<0.05, the difference between groups was statistically significant.

Previous reported studies and our results have presented evidence of vesicular internalization into recipient cells, however, the mechanism for the difference in SMC-MVs and Ost-MVs uptake remains elusive. SMC phenotypic switching, the shift of SMCs from a physiological contractile phenotype to a synthetic phenotype, is a pathological process that closely associates with atherogenesis and sequent calcification.^42, 43^ We questioned whether SMC phenotypic transition impacted on MVs internalization in vitro. Heparin or PDGF-BB was used to induce the contractile and synthetic phenotype of SMCs, respectively.^44, 45^ As shown in Figure 5C-H, SMCs treated with heparin led to upregulated expression of contractile markers α-SMA, SM22α, and CNN1(Calponin 1) and decreased expression of synthetic marker S100A4 (S100 calcium binding protein A4), indicating a contractile phenotype. Conversely, PDGF-BB treatment promoted a synthetic phenotype of SMCs with the loss of expression of α-SMA, SM22α and CNN1 and concomitant gain of S100A4 and MMP2 (Matrix metalloproteinase 2) expression.

Next, we assessed the calcification capacity of contractile or synthetic SMCs cultured in procalcific medium with an increased Ca^2+^ concentration. Elevated intracellular calcium was detected in synthetic SMCs in contrast to contractile cells after 48h (Figure 5I). This confirmed the linkage between calcification and SMC phenotypic switch. Then, flow cytometric and confocal microscopic analysis were performed to investigate the effects of different phenotypes of SMC on MVs uptake. Flow cytometric results revealed that contractile SMCs showed higher tendency to internalize SMC-MVs than Ost-MVs (98.5% *vs.* 49.2%), and the synthetic phenotype of SMCs facilitated the uptake of Ost-MVs compared with that of contractile SMCs (49.2% *vs.* 72.2%) (Figure 5J). Similar results were obtained by immunofluorescence (Figure 5K-L).

Taken together, these results demonstrated the endocytosis of Ost-MVs by recipient SMCs, and the internalization of Ost-MVs is regulated by a phenotypic shift from contractile SMCs to synthetic SMCs.

### 3.6 Both recruited Ost-MVs and resident SMC-MVs aggravated atherosclerotic calcification through the Ras-Raf-ERK pathway

Given that SMC-MVs and Ost-MVs not only act as well-established initial sites for mineral deposition, contain abundant calcification-related cargoes, we postulated that MVs released by mineralized cells may enhance the progression of calcification by delivering procalcific signals to recipient SMCs. Therefore, the effects of SMC-MVs and Ost-MVs on SMC mineralization were examined. As expected, using alizarin red S staining to visualize the formation of mineralized nodules, we found that the addition of either SMC-MVs or Ost-MVs significantly promoted calcium deposition and the formation of mineralized nodules (Figure 6A-B). Similarly, ALP activity in SMCs was increased in response to SMC-MVs and Ost-MVs, compared with that in the control group (Figure 6C).

**Figure 6.**
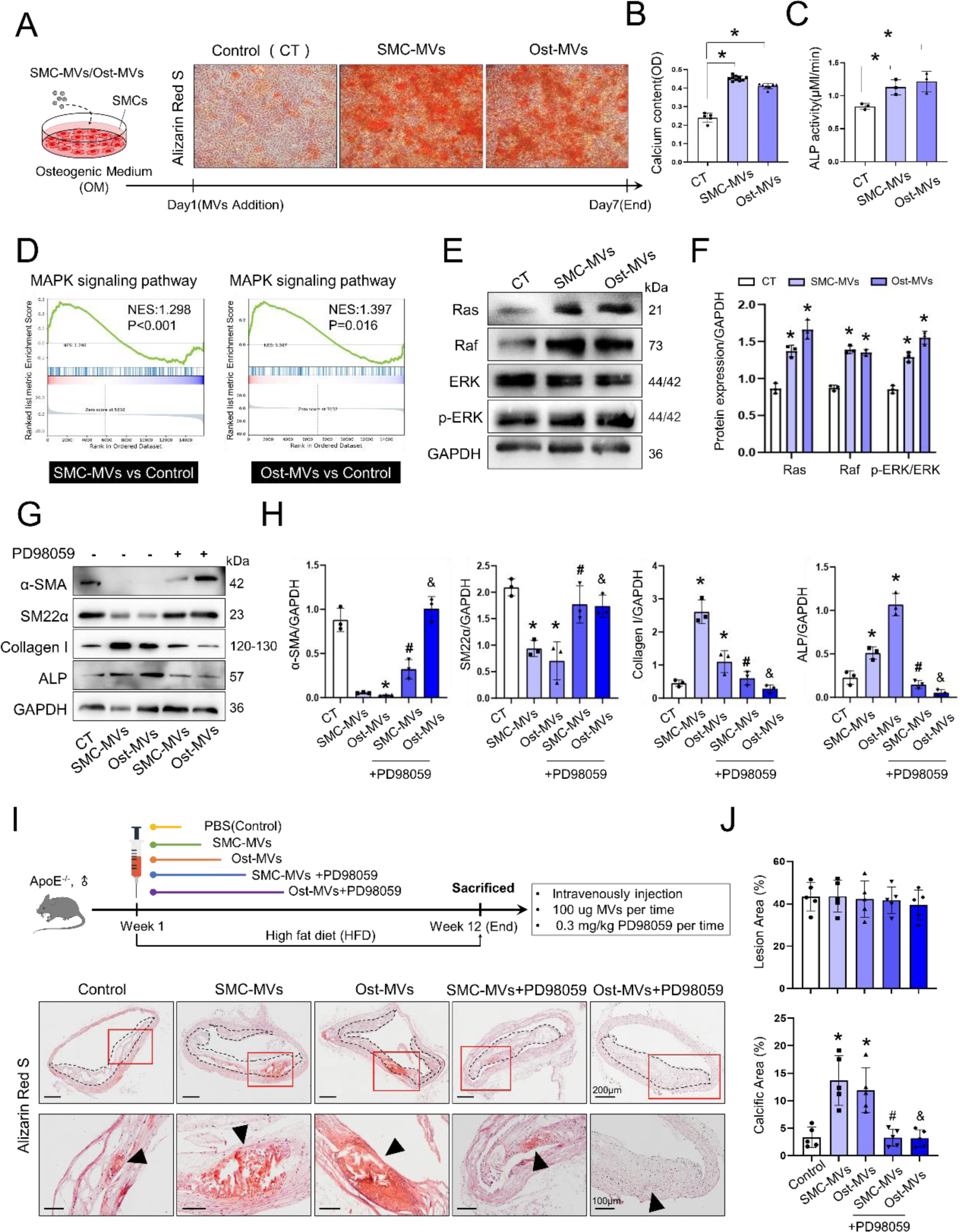
Both recruited Ost-MVs and resident SMC-MVs could expedite SMC osteogenic transdifferentiation and atherosclerotic calcification through the Ras-Raf-ERK signaling axis. SMC-MVs or Ost-MVs were co**-**cultured with SMCs in OM for 7d. (A) Representative images of in vitro SMC mineralization detected by Alizarin Red S staining. (B) Quantification of eluted calcium. (C) ALP activity of recipient SMCs at day 7. (D) GSEA results indicated hyperactivation of MAPK signaling in recipient SMCs after SMC-MVs or Ost-MVs treatment under OM condition compared with control group. NES, normalized enrichment score. (E-F) Western blotting and quantification of Ras, Raf, and ERK protein expression. (G-H) SMCs were pretreated with PD98059(10μ M) for 3d prior to the addition of SMC-MVs and Ost-MVs. Western blotting and quantification of SMC markers (α-SMA and SM22α) and osteogenic markers (Collagen І and ALP). (I) Experimental design of in vivo assay to detect the role of SMC-MVs and Ost-MVs in the development of atherosclerotic calcification and the role of MAPK signaling in this process. In vivo atherosclerotic calcification was detected by Alizarin Red S staining (n=5). Scale bar=200 μm. Designated regions indicated by red square frames are enlarged to show the details. Scale bar= 100 μm. (J and K) Quantification of the lesions for necrotic area and calcified area (n=5). *, P<0.05, the difference between SMC-MVs/Ost-MVs treatment and control groups was statistically significant. #, P<0.05, the difference between SMC-MVs and SMC-MVs+PD98059 groups was statistically significant. &, P<0.05, the difference between Ost-MVs and Ost-MVs+PD98059 groups was statistically significant.

To further study the mechanisms of MV–induced calcification, we performed RNA-seq to sequence the transcriptome profiles of SMCs in the control group, the SMC-MVs treatment group, and the Ost-MVs treatment group. As shown in Figure 6D, gene set enrichment analysis (GSEA) showed that MAPK signaling was positively enriched in the SMC-MVs treatment group (P < 0.001) and Ost-MVs treatment group (P = 0.016), compared with the control group, respectively. Indeed, the MAPK pathway has been reported to regulate the occurrence of vascular calcification.^46, 47^ Results of western blot confirmed the activation of the MAPK pathway after the stimulation of SMC-MVs or Ost-MVs, which was characterized by significantly upregulated expression of Ras, Raf, and p-ERK/ERK (Figure 6E-F). Notably, in different settings of calcification, the expression of osteogenic makers is classically associated with calcification.^48^ To explore the involvement of MAPK signaling in MV-mediated calcification, we pretreated SMCs with a ERK1/2 signaling inhibitor PD98059 prior to the addition of SMC-MVs and Ost-MVs. Subsequently, we assessed the expression of molecular makers of SMC osteogenic transdifferentiation in control group, SMC-MVs treatment group, Ost-MVs treatment group, SMC-MVs+PD98059 treatment group, and Ost-MVs+PD98059 treatment group, respectively. The addition of either SMC-MVs or Ost-MVs significantly decreased the gene and protein expression of typical SMC markers α-SMA and SM22α of recipient SMCs, and increased the expression of osteogenic makers collagen І and ALP at day 7. This pro-osteogenic effect of MVs was diminished by inhibition of ERK1/2 activity with PD98059 (Figure 6G-H, and Supplementary material online, Figure S6).

Finally, we investigated the role of SMC-MVs and Ost-MVs in the development of atherosclerotic calcification and the involvement of MAPK signaling in this process in vivo. ApoE^−/−^ mice on a high-fat diet were divided into control group, SMC-MVs treatment group, Ost-MVs treatment group, SMC-MVs + PD98059 treatment group, and Ost-MVs + PD98059 treatment group. After 12 weeks of high-fat diet, aorta tissue was selected for Alizarin red staining. Consistent with in vitro findings, the injection of either SMC-MVs or Ost-MVs significantly accelerated the progression of calcification. The treatment of PD98059 markedly abrogated the pro-mineralization effect of SMC-MVs and Ost-MVs, but PD98059 did not affect atherosclerotic lesion formation(Figure 6I-J).

In summary, these combined data indicated that both SMC-MVs and Ost-MVs accelerate calcification progression, and MV–induced calcification is regulated by the Ras-Raf-ERK axis.

## 4 Discussion

Vascular calcification portends worse clinical outcomes and predicts major adverse cardiovascular events. Currently, the key issue in the clinical management of vascular calcification is whether vascular calcification is reversible or amenable to therapy. The existence of calcification paradox implies that bone disorders have serious implications for cardiovascular health. Decreasing the efflux of calcium and phosphate from the bone can reduce the availability of calcium and phosphate as substrates for deposition of hydroxyapatite crystals in the arterial wall. Therefore, it is of great significance to find out a strategy to prevent or regress vascular calcification while the balance of bone–vascular axis is considered.

Intimal calcification is closely associated with overt atherosclerotic plaques. The mechanism by which calcification occurs resembles bone formation, but the environment of calcification is somewhat more similar to bone resorption.^49^ Substantial evidence from epidemiological research has revealed that low BMD is related to increased cardiovascular events, and worsened subclinical measures of vascular atherosclerosis. The Prospective Epidemiological Risk Factors Study Group, a 7.3-year study of follow-ups of postmenopausal men and women, showed that aortic calcification was associated with lower BMD, and bone loss, from the proximal femur.^50^ Likewise, the Multi-Ethnic Study of Atherosclerosis’s (MESA’s) Abdominal Aortic Calcium Study (AACS) showed that low BMD was strongly associated with increased deposition of coronary artery calcium.^51^ In the present study, we provided experimental data showing that hyperlipidemia-induced atherosclerotic calcification was accompanied by an obvious decrease in bone mass, suggesting the loss of bone mineral (Figure 1A, E-F). Adequate calcium and phosphorus supply is a prerequisite for both the occurrence of bone mineralization and vascular calcification. The mobilization of bone phosphorus and calcium into the blood following bone mineral loss may facilitate abnormal mineral deposition in vascular walls, leading to the initiation or worsening of vascular calcification.^21^

The calcification paradox suggests a link between bone and vasculature, and that there are possible mechanisms underlying the coincidence of bone loss and vascular calcification through the bone-vascular axis.^25, 52, 53^ The consensus among existing studies was that the MVs are the starting sites of mineralization, and the MV-mediated control of osteoblasts-and/or osteoclastogenesis is of particular relevance to vascular calcification.^54^ Hence, MVs may affect vascular health by acting as carriers that mediate the transport of calcium and phosphate from bone to circulation. In this study, both in vitro and in vivo results demonstrated that the occurrence of atherosclerotic calcification is accompanied with bone loss. This is characterized by a decrease in matrix-bound Ost-MVs and an increase in free circulating Ost-MVs (Figure 1B-D and G-I, and see Supplementary material online, Figure S2). Upon further investigation into the in vivo tracking of bone-released Ost-MVs, we observed their targeting of plaque lesions within the context of atherosclerosis. A strong fluorescent signal accumulation was evident in athero-susceptible regions such as the aortic arch and abdominal aorta (Figure 2). These findings indicates that Ost-MVs serve as mediators linking metabolic bone disorders with ectopic atherosclerotic calcification. However, the bone-resorbing osteoclasts^55^ and macrophages^56^ may also secrete functional matrix EVs and play roles in the calcification paradox, which still warrants future investigation.

SMC-MVs secreted from mineralized SMCs have been well-recognized as the main contributor to cardiovascular calcification.^57^ Interestingly, here we found that Ost-MVs could be mobilized from the bone into vascular wall, suggesting another mechanism responsible for pathologic calcification. To elucidate the cellular origin of MVs involved in atherosclerotic calcification, we initially conducted proteomic analysis on SMC-MVs and Ost-MVs, and further screened differentially expressed proteins between these two types of MVs. Our data showed that lysosomal membrane protein Lamp1 was specifically expressed in SMC-MVs, and Nid2 was expressed in Ost-MVs only (Figure 3A-E). This finding is consistent with a previous reported study, which also demonstrated that Lamp1 and Lamp2 were specifically expressed on calcifying SMC-derived EVs.^58^ Based on proteomic results, Lamp1 and Nid2 were used as biomarkers for SMC-MVs and Ost-MVs respectively in this study. We subsequently evidenced that extracellular, punctate deposits of SMC-MVs and Ost-MVs within calcific plaques, which was indicated by the co-localization of Lamp /Nid2 staining with ECM protein collagen І in the calcific plaque lesions (Figure 3F-H). These results demonstrated that both circulating Ost-MVs and resident SMC-MVs contribute to the formation of atherosclerotic calcification. Notably, although smaller types of vesicles are too small to be detected with regular light microscopy, these puncta that we showed might represent vesicle aggregates, as the aggregations has recently been proposed to promote calcification.^11^ Inconsistently, Chen et al. developed in vitro calcification models by high-phosphate treatment, and in vivo calcified mice by nephrectomy-induced chronic renal failure, cholecalciferol-overload, and periadventitially administering with CaCl_2_, they found a decrease in the Nid2 protein levels in calcified SMCs and aortas.^59^ Thus, the role of Nid2 used as biomarker for Ost-MVs is limited to the setting of HFD-induced atherosclerotic calcification in ApoE^−/−^ mice. And more specific molecular makers for Ost-MVs identification requires further investigation.

In this study, we further uncovered the mechanism for the recruitment of Ost-MVs into vascular matrix. Using transwell models combined with LPS treatment to induce vascular injury in the context of atherosclerotic calcification, we tested whether circulating Ost-MVs could be transported across the endothelial barrier. Our data indicated that LPS-induced endothelial inflammatory injury promoted transendothelial transport of Ost-MVs, whilst this was not observed under the condition of intact endothelium without LPS stimulation (Figure 4A-C). Of note, this study didn’t demonstrate the exact mechanism of MV transport. Except for direct transport through intercellular spaces, it has been reported that EVs internalized into recipient cells can also merge into endosomes, undergo transcytosis, move across recipient cells and be released into neighboring cells.^60^

Beyond that, the extracellular matrix change, mainly collagen І, is a histological feature shared by physiological bone mineralization and the pathological condition of atherosclerotic calcification, and significantly impacts the course of both processes.^11, 61, 62^ The remodeling of pro-calcifying collagen І, accompanied with the development of atherosclerotic lesion, may provide an explanation for the recruitment and deposition of circulating Ost-MVs to vascular matrix. By constructing collagen hydrogels to mimic the characteristics of ECM during atherosclerosis, we found that alternations in collagen І directed the accumulation and aggregation of Ost-MVs in the vascular matrix (Figure 4D-G). This was further confirmed by in vivo data. With the treatment of Halofuginone, a specific inhibitor of collagen I synthesis, the deposition of circulating Ost-MVs to the vascular matrix could be markedly abrogated (Figure 4H-K). However, the role of collagen in mediating SMC-MVs-induced calcification growth seems to differ from the finding of present study. Hutcheson et al demonstrated an inverse relationship between the collagen density and the volume of resulting mineral deposits in the extracellular space Using a ApoE^−/−^ mice and another strain of ApoE^−/−^ mouse deficient in matrix metalloproteinase 13 (ApoE^−/−^/MMP13^−/−^).^11^ This contradiction may be attributed to the limitations of in vitro assays to fully replicate the alternations of in vivo ECM environment during atherosclerosis.

In the progression of atherosclerotic calcification, collagen remodeling is a complex and dynamic process since collagen degradation and synthesis take place simultaneously within the same plaques.^63^ Moreover, current researches on the cell–matrix axis in calcifying EV release points towards an additional regulatory role of collagen on the cell level, activating feedback mechanisms to control quantitative EV release and the calcifying potential of EVs.^64^

Once EVs are released into the extracellular space, they undergo internalization.^65^ After uptake by recipient cells, the genetic materials and proteins are transferred from the vesicles to the recipient cells, which further regulate intracellular signaling pathways of the target cells.^65^ Here we also found that Ost-MVs, as well as SMC-MVs, could be incorporated in the recipient SMCs under pro-calcific conditions (Figure 5A-H). Interestingly, the uptake of MVs showed temporal differences in two types of MVs. Our subsequent study revealed that the regulation of this difference is attributed to SMC phenotypic transition. In detail, contractile SMCs tended to take up SMC-MVs rather than Ost-MVs, while the synthetic phenotype of SMCs facilitated the uptake of Ost-MVs (Figure 5J-L). Indeed, it has been reported that EV uptake is a highly context-specific process, which may depend on vesicular properties, the cell type and its pathophysiological state.^66^

In view of the significant roles of MVs in the formation and development of calcification, we proceeded with our investigation into the effects of recruited Ost-MVs and resident SMC-MVs on SMC mineralization progression and vascular calcification. Results from in vitro and in vivo experiments showed that both SMC-MVs and Ost-MVs promoted SMC osteogenic transdifferentiation and augmented the progression of calcification, which was mediated by Ras-Raf-ERK signal axis (Figure 6, and see Supplementary material online, Figure S6). A study on the signal transduction role of MVs performed by Chen et al., using co-cultured recipient normal VSMCs with cellular–derived MVs isolated from CKD rat VSMCs, demonstrated that MVs could be endocytosed by recipient cells, and facilitated the calcification of recipient VSMCs from normal rats, in concordance with our results. This effect, as reported by Chen et al., was partially attributed to the activation of both NOX and MAPK signaling, and they showed that inhibiting either pathway blocked the increase in intracellular calcium concentration, and decreased VSMC calcification.^67^ Another study using a diabetic atherosclerotic calcification model induced by Nε-Carboxymethyl-lysine (CML) in ApoE^−/−^ mice by Jing et al. found that, following the tail vein injection of SMCs-derived MVs, the aorta of the mice was obviously calcified.^68^ Of note, a limitation of our study in the assessment of osteogenic transdifferentiation is that we didn’t elucidate the potential candidates within MVs that functioning pro-calcifying effects. Accumulating evidence suggests that calcified MVs can induce expression of osteogenic marker genes in the target cells by delivery of dysfunctional miRNAs, thus influencing the phenotypic transition and function of recipient cells.^69^ Another limitation is that “dose-response” experiments were not performed to investigate the effects of Ost-MVs and SMC-MVs on vessel phenotypes in the context of atherosclerotic calcification. Indeed, there is a lack of evidence regarding the exact concentrations of Ost-MVs and SMC-MVs presented in the bone-vascular axis to support this study.

In conclusion, we presented an Ost-MVs-mediated mechanism to explain the calcification paradox and further enrich the knowledge of bone-vascular crosstalk (Figure 7). Since MVs function as central mediators in the process of bone formation and vascular calcification, targeted interventions that considering the bidirectional balance of the bone–vascular axis may provide novel therapeutic options for vascular calcification.

**Figure 7.**
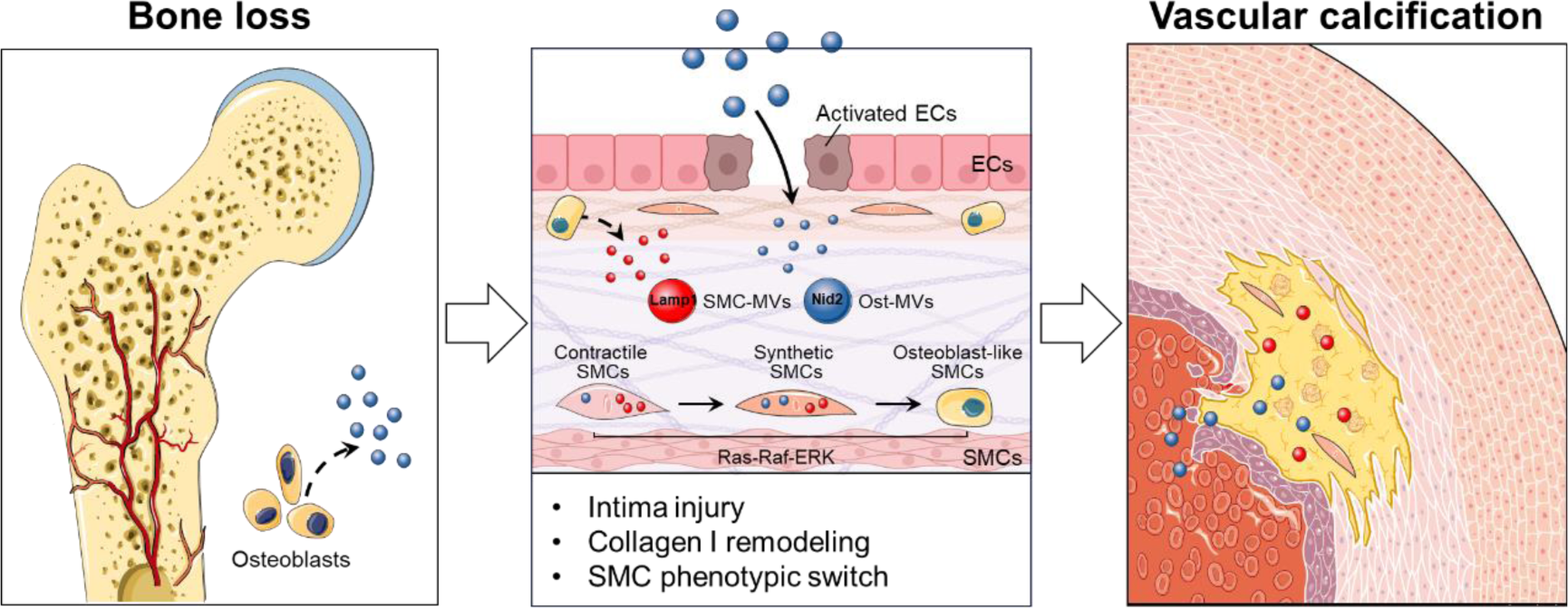
Ost-MVs released from bone during atherosclerotic calcification transport to plaque lesions and contribute to calcification progression. In the context of atherosclerotic calcification, Ost-MVs transport from bone to vascular matrix, and the underlying mechanisms involve vascular injury allowing Ost-MVs transendothelial transport, Collagen І remodeling-mediated Ost-MVs deposition, and SMC phenotypic switch to facilitate Ost-MVs uptake. Furthermore, recruited Ost-MVs and resident SMC-MVs promote osteogenic differentiation of recipient SMCs and aggravated calcification, depending on the Ras-Raf-ERK signaling pathway.

## Abbreviations

MVs: matrix vesicles
*SMCs*: *smooth muscle cells*
Ost-MVs: osteoblast-derived MVs
SMC-MVs: calcifying SMC-derived MVs
ApoE^−/−^ mice: apolipoprotein E-deficient mice
WT: wild-type
HFD: high-fat diet
ND: normal diet
BMD: bone mineral density
BV/TV: Bone volume/tissue volume
Tb.N: trabecular number
Tb.Th: trabecular thickness
Tb.Sp: trabecular separation
NM: normal medium
OM: osteogenic medium
ANX2/5/6: annexins 2, 5 and 6
Lamp1: Lysosome associated membrane protein 1
Nid2: Nidogen 2
LPS: lipopolysaccharide
α-SMA: α-smooth muscle actin
SM22α: smooth muscle 22α
CNN1: Calponin 1
S100A4: S100 calcium binding protein A4
MMP2: matrix metallopeptidase 2
ALP: alkaline phosphatase
MAPK: mitogen-activated protein kinase
ERK: extracellular regulated protein kinases
DiІ: 1,1’-dioctadecyl-3,3,3’,3’-tetramethylindocarbocyanine perchlorate
DiO: 3,3’-dioctadecyloxacarbocyanine perchlorate
PDGF-BB: platelet derived growth factor subunit B
NTA: nanoparticle tracking analysis
TEM: transmission electron microscope
GSEA: gene set enrichment analysis

## Acknowledgements

X.W., Y.S., and X.L. conceived and designed the experiments. X.W. and J.R. performed the experiments and wrote the manuscript, F.F., E.W., and J.L. analyzed and interpreted the data, W.H., Y.S., and X.L. reviewed the data and manuscript. All authors critically read and commented on the manuscript.

We thank Professor Lance L. Munn from Massachusetts General Hospital and Harvard Medical School (https://steelelabs.mgh.harvard.edu/), for editing the draft of this manuscript.

## Sources of Funding

This work was supported by the National Natural Science Foundation of China (Grant Nos.11932014 and 12372315, X.L.; 32071312, Y.S.; 82300505, X.W.) and Sichuan Science and Technology Program (Grant numbers 2022NSFSC0765 and 2022ZYD0079).

## Disclosures

The authors have declared that no competing interest exists.

## Notes

### Competing Interest Statement

The authors have declared no competing interest.

